# Transcriptional shut-off of MAP kinase signaling enables pluripotency maintenance during diapause

**DOI:** 10.1101/2023.12.17.572058

**Authors:** Tuo Zhang, Ryan J. Marina, Rab Prinjha, Karen Adelman, Alexander Tarakhovsky

## Abstract

Exposure of unicellular or multicellular organisms to adverse environmental conditions, including nutrient deprivation, may induce a state of suspended animation or diapause. The diapause minimizes the organism’s reliance on external energy sources and ensures survival. Among different forms of diapause, embryonic diapause, caused by a limited supply of nutrients to the growing embryos, is particularly challenging for the organism. Diapause embryos stop developing at the peak of pluripotent cell differentiation and maintain this undifferentiated yet entirely vital state despite the overall reduction in anabolic processes and genome-wide transcriptional repression. Using ES cells commonly employed to study the mechanism of embryonic diapause, we solve the paradox of the cell maintenance in an undifferentiated ES cell state during diapause. We find that broad transcriptional repression by long-term inhibition of the bromodomain and extra-terminal (BET) proteins causes diapause. These diapause ES cells upregulate a functionally linked group of genes encoding negative regulators of MAP kinase signaling (NRMKS), which play a crucial role in ES cell differentiation. We find that elevated NRMKS expression is a hallmark of the diapause cells and cells exposed to diapause-inducing conditions, including mTOR inhibition, and is required for the maintenance of ES cell pluripotency during diapause. Mechanistically, exposure of ES cells to diapause-inducing conditions leads to rapid decline of the Capicua transcriptional repressor (CIC) at the NRMKS gene promoters, followed by transcriptional upregulation of NRMKS genes. The mTOR and BET-dependent transcriptional switch supporting the undifferentiated state of the diapause ES cells suggests a broader usage of this mechanism in maintaining the undifferentiated state of metabolically dormant stem- or stem-like cells in different tissues.

## Main Text

The embryonic diapause is a discrete phenotypic state that occurs in response to a limited nutrient supply to newly conceived embryos in numerous species of animals, including mice(*1*). Diapause is characterized by reduced anabolism and diminished levels of various biosynthetic processes, including protein synthesis and gene expression(*2-5*). Remarkably, diapause-associated translational and transcriptional changes have no evident impact on the differentiation potential of the embryos, which upon restoration of the nutrient supply resume normal growth and give rise to healthy progeny(*1, 6*). The mechanism by which diapause embryos can maintain the undifferentiated potential for extended periods is unknown.

The diapause *in vivo* can be phenotypically imitated *in vitro* by nutrient deprivation of *ex vivo* isolated mouse blastocysts(*3*) or cultured embryonic stem (ES) cells(*3, 4*). The eukaryotic cell response to nutrient supply is governed by the mTORC1/2 protein complexes(*7*). Accordingly, pharmacological suppression of mTOR imitates nutrient deprivation and causes a diapause-like state *in vitro*(*3, 4*). The diapause-like condition in ES cells can also be triggered by *Myc* family protein deficiency(*5*). The latter finding appears paradoxical given the crucial role of *Myc* in regulating both the expression of genes that govern ES cell growth and genes that control pluripotency(*8, 9*). While the reduced metabolic and growth rates of *Myc*-deficient ES cells have a straightforward mechanistic explanation, it remains unclear how *Myc*-deficient ES cells with a broad transcriptional repression maintain a naïve cell state.

We hypothesized that maintenance of ES cell pluripotency during diapause relies on a mechanism that couples the diapause-promoting signals with processes that support the ES cell undifferentiated state. To address the nature of this potential mechanism, we created conditions of a broad and long-lasting, yet reversible, transcriptional repression by culturing mouse ES cells in the presence of I-BET151, the pharmacological inhibitor of the bromodomain-containing BET proteins(*10*). The BET proteins BRD2, BRD3, and most prominently, BRD4 contribute to gene expression by linking the acetylated histones and, to a lesser extent, other nuclear proteins that control the RNA Pol II activity(*11, 12*).

The exposure of ES cells to incrementally increased I-BET concentration results in the generation of ES cells, hereafter defined as I-BET resistant (I-BETR), that can exist in the presence of I-BET at a concentration prohibitive for the bromodomain-dependent chromatin binding(*13*) (Fig. S1A). The I-BETR ES cells express the naïve ES cell marker alkaline phosphatase (AP)(*14*), are smaller and grow at considerably slower rate than the control ES cells. (Fig. 1A-B). The latter feature of I-BETR ES cells explains the greatly reduced size of I-BETR ES cell colonies that have otherwise a dome-like shape typical for control ES cells (Fig. 1A) The I-BETR cells have decreased biosynthetic activity, as indicated by reduced overall and *de novo* RNA and protein synthesis (Fig. 1D). The metabolic state of I-BETR ES cells is characterized by reduced levels of oxidative phosphorylation as judged by lower basal and maximal oxygen consumption rates (OCR) (Fig. 1C). The declined oxidative phosphorylation capacity is coupled with reduced levels of glycolysis as judged by reduced basal and maximal extracellular acidification rates (ECAR) before and after adding the mitochondrial electron transport chain inhibitors (Fig. 1C). The described features of I-BETR ES cells are consistent with the previously described phenotypes of ES cells that were rendered diapause by mTOR inhibition(*3*), nutrient deprivation(*4*), or combined *Myc*-family protein deficiency(*5*). Several lines of evidence suggest that acquiring the diapause-like state by I-BETR ES cells reflects cell adaptation rather than a selection of the rare ES cell variant. First, the progression towards the high concentration of I-BET was associated with minimal cell attrition at each step of increasing I-BET concentration (Fig. S1A-B). Second, removal of I-BET fully and quickly reverses the diapause-like phenotype of I-BETR ES cells (Fig. 1B-C). Finally, after a short-term (12-14 hours) removal of I-BET, the I-BETR ES cells could generate chimeras upon injection into the C57BL/6J mice-derived blastocysts (Fig. 1E). This finding underscores the unaltered pluripotent capacity of I-BETR ES cells and is consistent with the reversibility of diapause(*3, 5*).

The long-term inhibition of BET proteins is associated with significant gene expression changes in I-BETR ES cells. The overall number of the affected genes not only underscores the breadth and scope of the transcriptional impact of chronic I-BET exposure, but also highlights cellular processes and biochemical pathways that can be linked to diapause(*2-5*) (Fig. 2A-B, Table S1). Certain functionally linked gene groups such as ribosomal protein genes encoding components of the large and small ribosome subunits as well as mitochondria-localized ribosomes are suppressed in I-BETR ES cells (Fig. 2C, Table S1). The downregulation of ribosomal protein genes correlates with the reduction in cellular RNA levels as well as *de novo* RNA synthesis (Fig. 1D). Given the overwhelming contribution of the ribosomal RNA to the overall cellular RNA content(*15, 16*), the reduced levels of global RNA are reflected by the large changes in ribosomal RNA expression. A combination of repressed ribosomal RNA and ribosomal protein expression is likely to be one of the major causes of the reduced protein synthesis in the I-BETR ES cells (Fig. 1D). Similar to the ribosomal protein gene expression, many of the transcriptional changes in the I-BETR cells can be logically linked to the known diapause-associated anabolic or catabolic changes (Table S1). We also noticed an upregulation of genes encoding negative regulators of protein kinase activity and signaling (Fig. 2A). The latter observation suggested that broad transcriptional inhibition by I-BET leads to suppression of signaling events that promote cell growth while preserving the pluripotent state of I-BETR ES cells by antagonizing differentiation-inducing signals.

The possibility of signaling control of the pluripotent state of I-BETR ES cells became apparent after comparing the impact of long-lasting I-BET suppression on genes that control ES cell pluripotency and differentiation (Fig. 2D-E). A closer look at the pattern of these gene expression changes shows a much more severe impact of long-lasting BET inhibition on differentiation-inducing genes than the genes control ES cell pluripotency. Only a few pluripotency genes, such as *Prdm14, Nanog* and *Sall1* are markedly downregulated in the I-BETR ES cells (Fig. 2D-E), while the rest of the critical pluripotency genes are expressed at control-like levels (Fig. 2D-E). In contrast to the pluripotency-controlling genes, the expression of numerous genes that drive ES cell differentiation into various lineages is nearly extinguished in the I-BETR ES cells (Fig. 2D-E). Remarkably, the expression of differentiation-promoting genes in the I-BETR ES cells was as low as in ES cells treated with the mix of GSK3β and MAP kinase inhibitors (2i)(*14*) (Fig. 2E). The MAP kinase signaling pathway plays a defining role in ES cell pluripotency and differentiation(*17*). Pharmacological inhibitors of MAP kinase activity block ES cell differentiation and enable highly pluripotent ES cell maintenance(*14*). The striking similarity between the impact of 2i and the long-lasting BET inhibition suggested a possible key role of MAP kinase signaling inhibition in the maintenance of naïve phenotype of the I-BETR ES cells.

Indeed, a comparative analysis of gene expression patterns between I-BETR ES cells and previously described *Myc*-deficient ES cells(*5*) revealed negative regulators of MAP kinase signaling pathway *Dusp4, Dusp6* and *Spry4(18, 19)* among a small group of commonly affected genes. Additionally, *Dusp4, Dusp6* and *Spry4* are among the 5 upregulated genes that are shared by I-BETR ES cells and nutrient deprived ES cells, or ES cells treated with mTORC1/2 inhibitor (Fig. 3A). An increase in *Dusp4, Dusp6* and *Spry4* mRNA expression levels in the I-BETR ES cells is accompanied by a rise of the corresponding protein levels and a concomitant decline of phosphorylated ERK (pERK) levels (Fig. 3B-C). Finally, the siRNA-mediated knockdown of *Dusp4, Dusp6* or *Spry4* or the combined knockdown of all three genes led to the differentiation of I-BETR ES cells, but did not substantially affect control ES cells, as measured by alkaline phosphatase expression and changes in ES cell morphology (Fig. 3D).

The described transcriptional activation of negative regulators of MAP kinase signaling (NRMKS) is among the earliest events that occur in response to mTOR inhibition in ES cells. Indeed, inhibition of mTOR affects the transcription of hundreds of genes within minutes, as determined by measurements of newly synthesized RNA using Transient Transcriptome (TT)-seq(*20, 21*) (Fig. 4A, 614 genes upregulated, 451 downregulated, Table S2). The rapidly activated and suppressed genes are confirmed by RNA-seq and contribute to functionally opposing catabolic or anabolic processes (Fig. 4B). The rapid speed and the breadth of transcriptional changes caused by mTOR suppression raises questions about the nature of transcriptional regulators that operate downstream of mTOR and relay the signaling state of mTOR to genes that contribute to cell adaptation to nutrient deprivation. Remarkably, the NRMKS genes are part of the non-catabolic genes that form a functionally coherent group that become activated in response to mTOR suppression (Fig. 4A, C-D). This group includes dual-specificity protein phosphatase *Dusp6, Dusp4* and *Dusp1* as well as members of Sprouty family signaling antagonist *Spry4, Spry2* and *Spry1* (Table S2). The upregulation of NRMKS in response to mTOR inhibition is an event of a major significance with respect to cell maintenance at the pluripotent state during nutrient deprivation. In ES cells or cells of other types, NRMKS support the signal-induced transcriptional negative feedback mechanism of MAP kinase signaling(*18, 19*). The signal-induced activation of MAP kinase leads to dissociation of the Capicua transcriptional repressor (CIC) from NRMKS gene promoters followed by NRMKS upregulation and inactivation of MAP kinase signaling(*22*). Our data show that a similar mechanism is involved in suppression of MAP kinase signaling in response to the diapause-induced signals. Pharmacological inhibition of mTORC1/2 in ES cells triggers a major decline in CIC occupancy of the NRMKS promoters that coincides chronologically with the up-regulation of NRMKS genes (Fig. 4E-F). These data suggest CIC dissociation from the NRMKS promoters as a key transcriptional event that prevents differentiation under conditions that instigate diapause.

The described mTOR and BET controlled mechanism of diapause ES cell maintenance in an undifferentiated state has several implications. Our studies explain the high efficiency and pluripotency potential of ES cells derived from the diapause embryos(*23, 24*). Pharmacological inhibition of MAP kinase is commonly used for maximizing the ES cell pluripotency by blocking ES cells in an undifferentiated state(*14, 25*). The ability of BET inhibitor to emulate the effect of MAP kinase signaling inhibitors via transcription-dependent mechanism points to the possible usage of BET inhibitors for pluripotency preservation in mouse and possibly human ES cells as well as *in vitro* maintenance of early embryos. Finally, the diapause induction by blockade of bromodomain-mediated BET interaction with chromatin suggests that exogenous factors such as pathogen-derived proteins that mimic the BET-recognized histone sequences and act as BET antagonists(*26, 27*)can be responsible for triggering the diapause-like metabolic state in the virus-infected cells(*28-30*).

