## Supplementary material for "Transcriptional shut-off of MAP kinase signaling enables pluripotency maintenance during diapause": Figures 1-4

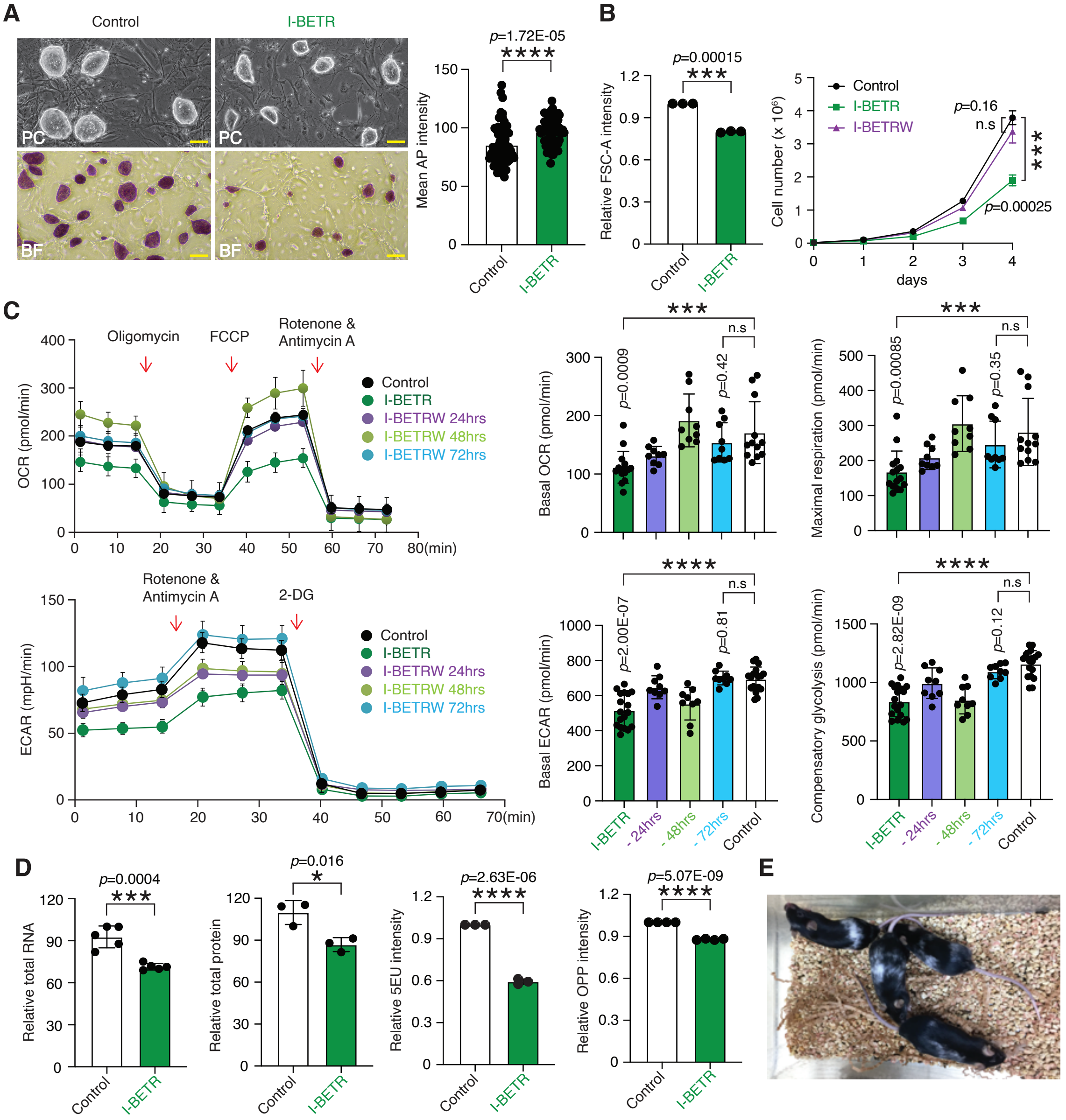


**Fig. 1. Long-term inhibition of bromodomain-dependent BET protein binding triggers the diapause-like state in ES cells.** (**A**), The I-BETR ES cells form dome-shape colonies and express alkaline phosphatase (AP) at control levels. The colonies of control and I-BETR ES cells are shown (left panel). PC, phase contrast microscopy; BF, bright field microscopy. Scale bar, 100µm. Bar graph (right panel) represents comparable expression levels of AP in control and I-BETR ES cells. Values represent normalized mean ± SD. Control ES cells, n=65; I-BETR ES cells, n=71. ****, $p<0.0001$, Unpaired t-test. (**B**), The size (left panel) and the growth rates (right panel) of control and I-BETR ES cells are shown. The size of ES cells was measured by flow cytometry assay and the data was quantified after elimination of Tdtomato+ feeder cells. The withdrawal of I-BET (I-BETRW) restores the I-BETR ES cell growth to control levels. Values represent normalized mean ± SD, n=3. n.s, no significance, ***, $p<0.001$, Unpaired t-test. (**C**), Altered metabolism in I-BETR ES cells. The oxygen consumption rate (OCR) (upper panel) and extracellular acidification rate

(ECAR) were quantified by Seahorse assay. The withdrawal of I-BET (I-BETRW) restores the I-BETR ES cell metabolism to control levels. Values represent normalized mean ± SD, n=6-12. n.s, no significance, ***, $p<0.001$, ****, $p<0.0001$, Unpaired t-test. (**D**), Reduced total amount (left two panels) and rates of de novo RNA and protein synthesis (right two panels) in I-BETR ES cells. The rates of de novo RNA and protein synthesis were measured by click chemistry assays with 5EU (5-Ethynyl-Uridine) labeling for nascent transcribed RNA and OPP (O-Propargyl-Puromycin) labeling for de novo translated protein. Values represent normalized mean ± SD, n=3-4. *, $p<0.05$, ***, $p<0.001$, ****, $p<0.0001$, Unpaired t-test. (**E**), Generation of chimeras by I-BETR ES cells. The I-BETR were incubated in I-BET-free medium for 12-14 hours and injected into the C57BL/6J blastocysts. White coat color indicates the degree of chimerism.

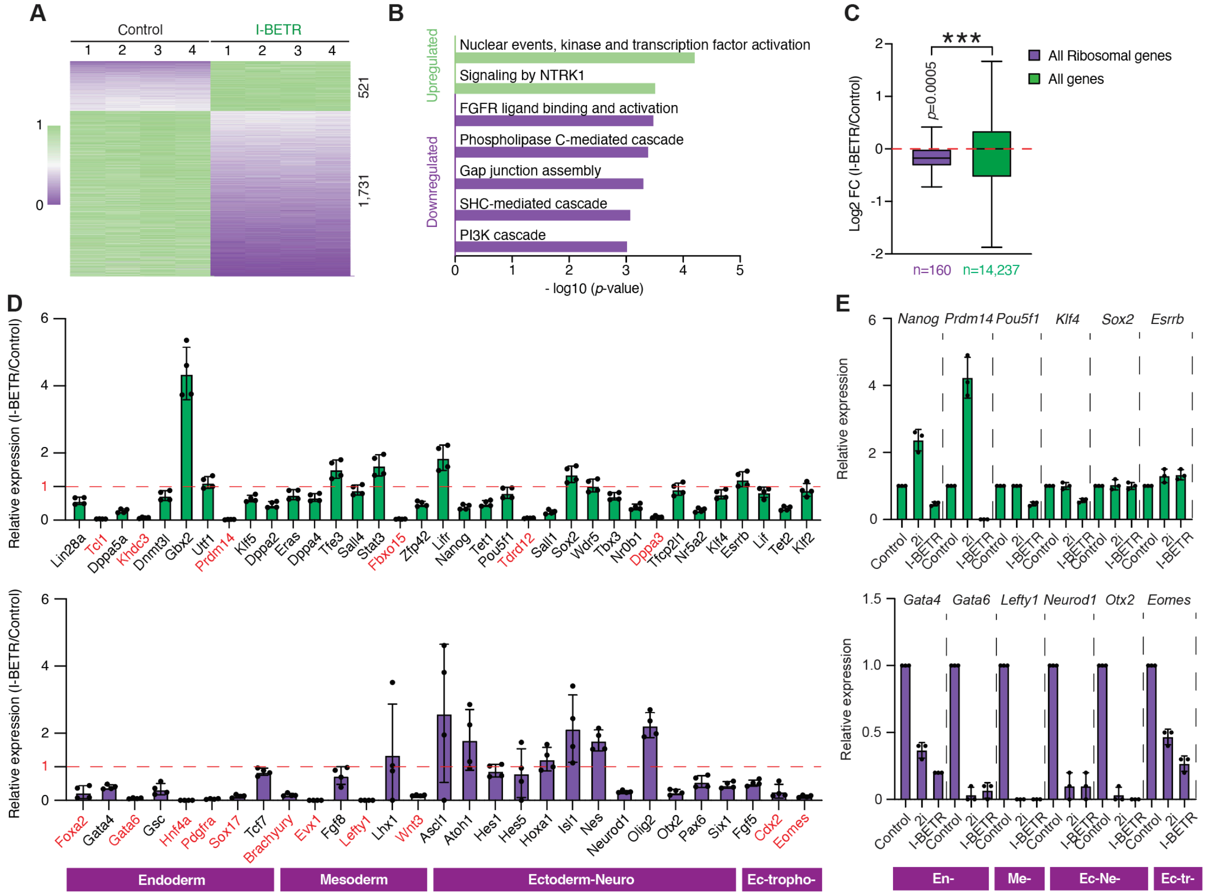


**Fig. 2. Transcriptional reprogramming of I-BETR ES cells is associated with repression of differentiation-inducing genes.** (**A**), The total RNA expression pattern in I-BETR ES cells. Heatmap shows 1,731 downregulated and 521 upregulated genes in I-BETR versus control ES cells. Each condition is represented by independent biological replicates (n=4). Scale bars represent relative expression normalized to max expression values determined by DESeq2-calculated normalized read counts. (**B**), The Reactome pathways analysis based on the top 200 upregulated or downregulated genes with adjust *p*-value < 0.0001 in I-BETR ES cells. The bar plots show - log10 (*p*-value). (**C**), Correlation analysis of all the ribosomal genes and global gene expression levels in I-BETR versus Control ES cells. Values represent log2 Fold Change (FC). ***, *p*<0.001, Mann-Whitney U test. (**D**), Expression of key pluripotency- (upper panel) and differentiation-specific (lower panel) genes in I-BETR ES cells. The bar plots represent relative expression of indicated genes in I-BETR compared to control ES cells based on the RNA-seq analysis, n=4. The genes display drastic downregulation are highlighted in red. (**E**), Transcriptional suppression of differentiation-specific genes in I-BETR ES cells. The comparative Q-PCR analysis of pluripotency-specific (upper panel) and differentiation-specific (lower panel) genes in control and I-BETR ES cells. 2i treated ES cells serve as the naive control for the assay. Values represent normalized mean ± SD, n=3.

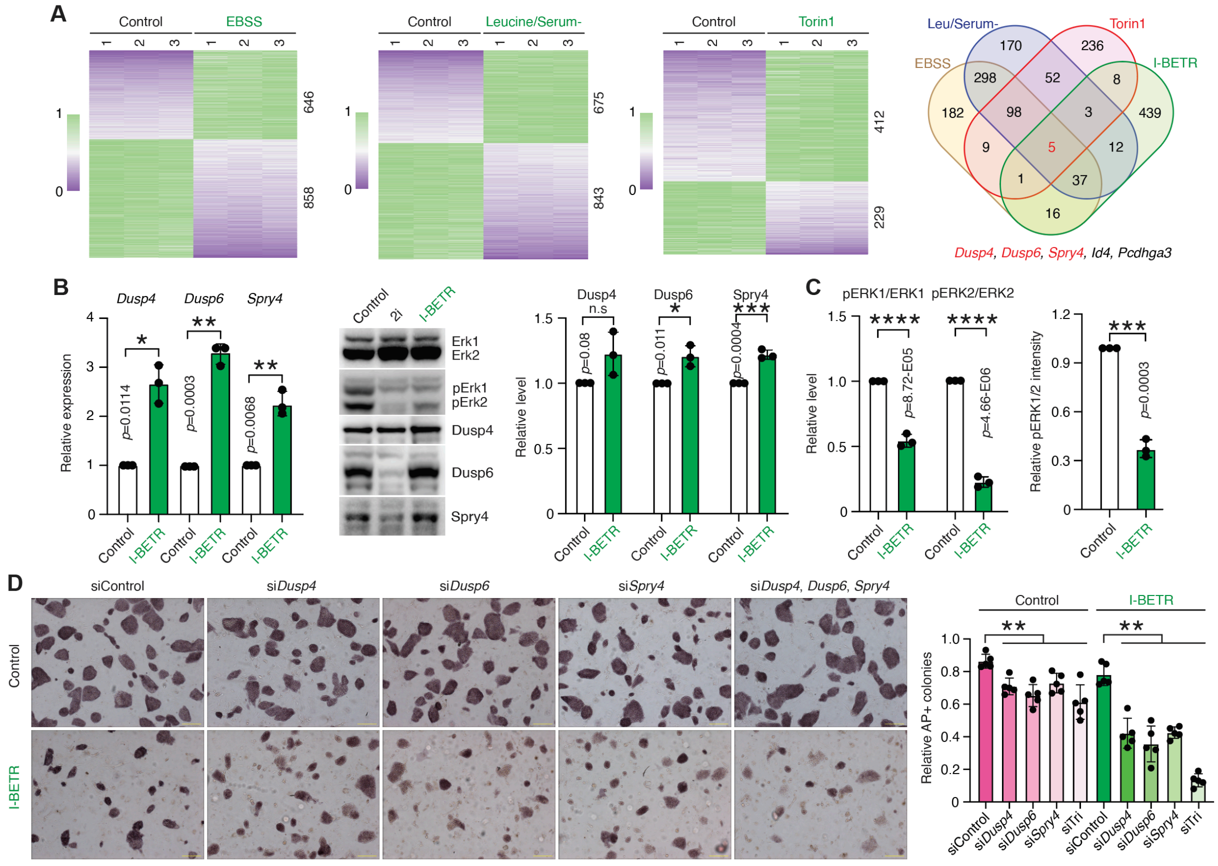


**Fig. 3. The transcriptional activation of negative regulators of MAP kinase in I-BETR ES cells.** (**A**), The negative regulators of MAP kinase are among genes that are shared by I-BETR ES cells and nutrient-deprived ES cells. Heatmaps show differentially expressed genes in EBSS starved leucine/serum deprived and Torin1 treated ES cells (left three panels). Each condition is represented by independent biological replicates (n=3). Scale bars represent relative expression normalized to max expression values determined by DESeq2-calculated normalized read counts. Venn diagram (right panel) represents overlapping up-regulated genes in all the tested conditions, including I-BETR ES cells. (**B**), Expression of *Dusp4*, *Dusp6* and *Spry4* mRNA (left panel) and protein (middle and right panel) in I-BETR and control ES cells. The mRNA and protein expression were quantified by Q-PCR and Western blotting respectively. Error bars indicate SD, n=3. n.s, no significance, *, $p$<0.05, **, $p$<0.01, ***, $p$<0.001, Unpaired t-test. (**C**), Reduced activity of ERK1/2 and pERK1/2 in I-BETR ES cells. The Erk activity was quantified by measuring the levels of the phospho-Erk by Western blotting (left panel) or immunofluorescence (right panel). Values represent relative mean ± SD, n=3. ***, $p$<0.001, ****, $p$<0.0001, Unpaired t-test. The impact of I-BETR on Erk activity was also compared to the impact of "2i" inhibitors shown by Western blotting in 3b (middle panel). (**D**), The siRNA mediated suppression of the indicated genes leads to I-BETR ES cells differentiation and reduction in number of colonies. The differentiation state of siRNA untreated or treated control or I-BETR ES cells was accessed by measuring the alkaline phosphatase (AP) expression. Scale bar, 100μm. The bar diagram shows the number of AP positive ES colonies. Values represent relative mean of undifferentiated colonies (16-44) ± SD, n=5. **, $p$<0.01, Unpaired t-test.

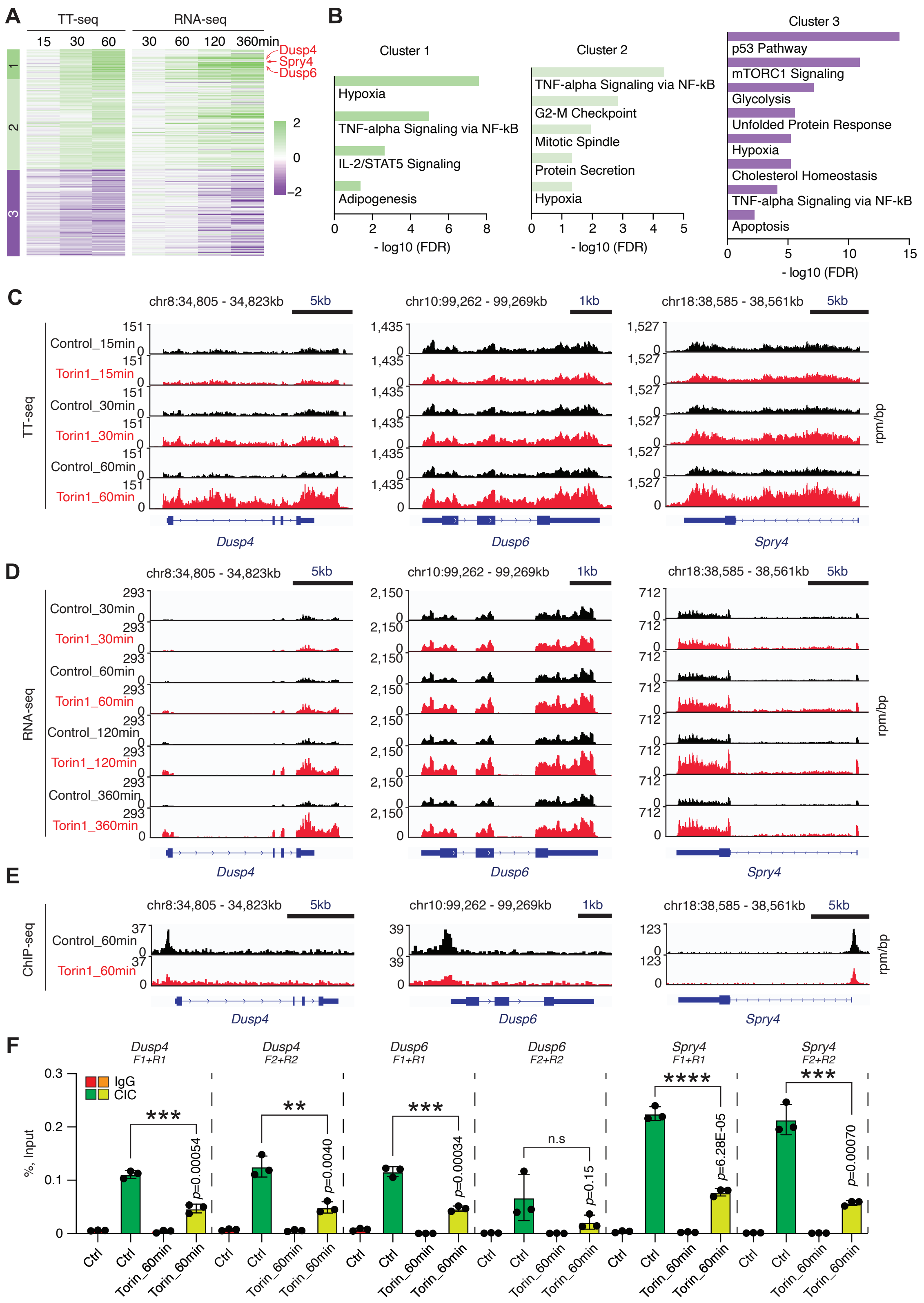


**Fig. 4. Rapid transcriptional activation of gene encoding negative regulators of MAP kinase in response to pharmacological mTOR inhibition.** (A), *Dusp4*, *Dusp6* and *Spry4* are part of the

rapid gene transcriptional response program triggered by mTOR inhibition. The impact of mTOR inhibition on gene transcription has been measured by nascent RNA sequencing (TT-seq) (left panel) and compared to the expression pattern of mature RNA sequencing (RNA-seq) (right panel). The gene expression changes are displayed as a heatmap with 3 clusters. The arrows indicate the positions of *Dusp4*, *Dusp6* and *Spry4* in the heatmap. (**B**), MSigDB gene ontology analysis based on the genes display nascent expression changes in each cluster shown in a. The representing pathways are the top hits with FDR adjusted *p*-values < 0.05. The bar plots show -log10 (FDR) values. Nascent (**C**) or mature (**D**) mRNA expression levels of *Dusp4*, *Dusp6* and *Spry4* at the indicated times of Torin1 treatment are shown in browser shots. CIC rapidly dissociates from promoters of *Dusp4*, *Dusp6* and *Spry4* upon mTOR inhibition in ES cells. (**E**), Browser shots represents CIC signals at *Dusp4*, *Dusp6* and *Spry4* gene loci in control and Torin1 treated ES cells for 60min. rpm/bp, reads per million per base pair. (**F**), ChIP-qPCR analysis represents CIC binding patterns at promoters of *Dusp4*, *Dusp6* and *Spry4* in control and Torin1 treated ES cells for 60min. Values represent normalized mean ± SD, n=3. *, $p<0.05$, ***, $p<0.001$, ****, $p<0.0001$, Unpaired t-test.
