## Supplemental Material and Data for "Transcriptional shut-off of MAP kinase signaling enables pluripotency maintenance during diapause"

### Materials and Methods

#### ES cell culture

The parental "CY2.4" ES line was derived from tyrosinase$^{-/-}$ (white, albino) homozygous mouse embryos of B6(Cg)-Tyrc-2J/J C57BL/6J background(*1*). ES cells were constantly maintained on the inactivated mouse embryonic fibroblasts (MEFs) by the treatment with 10 μg/ml final concentration of mitomycin C (M4287, Sigma-Aldrich) at 37°C with 5% $CO_2$ for 2.5-3.0 hours. Parental ES cells were cultured in ES medium: 40% EmbryoMax DMEM (SLM-220, Millipore), 40% KnockOut DMEM (10829018, Thermo Fisher Scientific), 7.5% FBS (100-500, Gemini), 7.5% KnockOut Serum Replacement (10828028, Thermo Fisher Scientific), 0.1 mM EmbryoMax MEM non-essential amino acids (TMS-001-C, Millipore), 1 × EmbryoMax Nucleosides (ES-008-D, Millipore), 50 units/ml EmbryoMax penicillin/streptomycin (TMS-AB2-C, Millipore), 0.1 mM EmbryoMax 2-Mercaptoethanol (ES-007-E, Millipore), 2 mM L-Glutamine (TMS-002-C, Millipore) and 1,000 units/ml LIF (ESG1107, Millipore). 2i ES cells were cultured in 2i medium: 50% DMEM/F-12 (11320033, Thermo Fisher Scientific), 50% Neurobasal medium (21103049, Thermo Fisher Scientific), 1 × B27 supplements (17504044, Thermo Fisher Scientific), 1 × N2 supplements (17502048, Thermo Fisher Scientific), 0.1 mM EmbryoMax MEM non-essential amino acids (TMS-001-C, Millipore), 50 units/ml EmbryoMax penicillin/streptomycin (TMS-AB2-C, Millipore), 2 mM L-Glutamine (TMS-002-C, Millipore), 1% KnockOut Serum Replacement (10828028, Thermo Fisher Scientific), 10 mM 1-Thioglycerol (M6145, Sigma-Aldrich), 1 μM PD0325901 (72182, Stemcell Technologies), 3 μM CHIR99021 (72052, Stemcell Technologies), and 1,000 units/ml LIF (ESG1107, Millipore). I-BETR ES cells were cultured in ES medium containing 4 μM I-BET151 (GSK1210151A). Earle's Balanced Salt Solution (EBSS) (24010043, Thermo Fisher Scientific) and leucine-free DMEM (226-024, Crystalgen) without serum were used for cell starvation assays. To inhibit the mTOR signaling, ES cells were maintained in ES medium containing 1μM Troin1 (S2827, Selleckchem).

#### Generation of I-BET resistant (I-BETR) ES cells and chimerism.

Parental CY2.4 albino B6 ES cells derived from B6(Cg)-Tyrc-2J/J embryos were constantly maintained on the mitomycin treated mouse embryonic fibroblasts (MEFs). Parental ES cells were rendered to I-BET resistant (I-BETR) with incrementally increase the concentration of I-BET151, starting from 250nM and ending with 4,000nM (4μM) (Fig. S1). The I-BETR ES cells were maintained one passage after removal of MEFs on the 0.5% gelatin treated plate before chimerism assay. After 12-14 hours of withdraw of I-BET from the culture medium (described before), the I-BETR ES cells were used for morula aggregations.

In brief, C57BL/6J inbred stock was used as embryo donors for aggregation with CY2.4 albino I-BETR ES cells and as pseudo pregnant surrogates. Embryos were collected at E2.5 from super-ovulated C57BL/6J females. Zona pellucida of embryos were removed by using acid Tyrode's solution (T1788, Sigma). I-BETR ES cell colonies were treated with 0.05% Trypsin-EDTA for getting single cell suspension and 10-20 cells was aggregated with each zona-free embryo. Recombined chimeric embryos were cultured overnight in micro drops of EmbryoMax advanced

KSOM embryo medium (MR-101-D, Sigma) covered with mineral oil (9305, Fujifilm) at 37 °C in 6% CO2. In the next morning, morulae and/or blastocysts were transferred into the uteri of pseudo pregnant E2.5 C57BL/6J surrogates. Chimerism was judged at birth by the presence of black eyes and later by the coat pigmentation.

**Western blotting**

$1 \times 10^6$ cells were collected and lysed on ice for 1 hour in 100 µl low salt (150mM NaCl) RIPA buffer (89900, Thermo Fisher Scientific) containing 2 × Protease Inhibitor Cocktail (P8340, Sigma-Aldrich), 1 × Phosphatase Inhibitor Cocktail Set I (Millipore, 524624), 331nM TSA (1406, Tocris) and 2µl of Benzonase Endonuclease (70664, Millipore) supplemented with 2mM $MgCl_2$.. 7.7 µl of 5M NaCl was add for every 100 µl of RIPA buffer to increase the salt concentration to 500mM. Cell lysate was then incubated on a spinning wheel in the cold room (4°C) for 1 hour, following with spinning down at 12,000g for 15min to separate the supernatant as the whole cell lysate. For western blotting, the whole cell lysate was mixed with 1 × NuPAGE LDS Sample Buffer (NP0007, Thermo Fisher Scientific) and 1 × NuPAGE Sample Reducing Agent (NP0004, Thermo Fisher Scientific) and denatured at 70°C for 10 minutes. 50,000 cells derived lysate was loaded into 12% Bis-Tris Protein gels (NP0341BOX, Thermo Fisher Scientific) and separated at 150 volts for 45-50 minutes. Protein gels then transferred onto Trans-Blot Turbo Mini 0.2 µm nitrocellulose membranes (1704158, Bio-Rad) and stained with ponceau S Staining Solution (A40000279, Thermo Fisher Scientific) to ensure equal transfer. Membranes were sequentially incubated with primary antibodies and secondary antibodies conjugated with horseradish peroxidase antibodies. After Incubation with SuperSignal West Dura Extended Duration Substrate (34075, Thermo Fisher Scientific), membranes were scanned by using ChemiDoc Imaging System (12003153, Bio-Rad) and the data was analyzed by Image Lab (Version 6.1). Western blotting for the investigation of pERK1/ERK2 and pERK1/ERK2 levels in the cells were performed by using stripping method. Membranes for pERK1/ERK2 detection was stripped with Restore PLUS Western Blot Stripping Buffer (46430, Thermo Fisher Scientific) for 15min at room temperature and blocked for 3 hours with 5% BSA TBST. Membranes were then reused for pERK1/ERK2 detection. Antibodies used in the study including Anti-pERK1/ERK2 (9101S, Cell Signaling), Anti-ERK1/ERK2 (9102S, Cell Signaling), Anti-Dusp4 (ab216576, Abcam), Anti-Dusp6 (ab76310, Abcam) and Anti-Spry4 (ab228712, Abcam).

**Analysis of nascent transcription or *de-novo* protein synthesis**

Total nascent transcription (5-Ethynyl Uridine, 5-EU) or de novo translation (O-propargyl-puromycin, OPP) were assessed in ES cells using Click-iT RNA Alexa Fluor 488 Imaging Kit and Click-iT Plus OPP Alexa Fluor 647 Kit respectively according to the manufacturer's instructions (C10329 and C10458, Thermo Fisher Scientific). For analysis nascent RNA synthesis, ES cells were incubated for 1 hour in ES medium supplemented with 0.5 mM 5-EU. To measure de novo protein synthesis, ES cells were incubated for 30 minutes in ES medium supplemented with 10 µM OPP. Cells treated with 1mg/ml of Actinomycin D (15021, Cell Signaling) for 2 hours and 50µg/ml of Cycloheximide (239763, Millipore) for 4 hours were used as the negative controls for the assays. The collected cells were fixed for 15 min at room temperature in PBS supplemented with 3.7% paraformaldehyde (19208, Electron Microscopy Sciences) following by permeabilized in PBS supplemented with 1% BSA (A7906, Sigma-Aldrich) and 0.1% Saponin (SAE0073,

Sigma-Aldrich) for 10 min at room temperature. The copper(I)-catalyzed alkyne-azide cycloaddition (CuAAC) was performed by using the indicated kits respectively. Samples were analyzed on the BD LSRII flow cytometry. Data were analyzed using the FlowJo software (Tree Star, Ashland, OR) and showed similar variance.

**Analysis of intracellular pERK1/2 level**

$1 \times 10^6$ ES cells were collected and pelleted by spinning for 3min at 300g. Cells were resuspended in 1 ml 1 × DPBS (14190144, Thermo Fisher Scientific). Formaldehyde (28908, Thermo Fisher Scientific) was added to obtain a final concentration of 4% and the cells were fixed for 10 min at room temperature. Cells were then chilled on ice for 2min and washed twice with 1 ml 1× DPBS. Cells next were permeabilized by resuspending in 200 µl of 0.1% Saponin (SAE0073, Sigma-Aldrich) in 3% BSA DPBS and incubated for 15 min at room temperature. Permeabilization buffer was discarded, and cells were resuspended in 100 µl of 0.1% Saponin in 3% BSA DPBS containing 1:200 diluted Anti-pERK1/ERK2 (9101S, Cell Signaling), and incubated for 30 min at room temperature. Primary antibody was discarded, and cells were resuspended in 100 µl of 0.1% Saponin in 3% BSA DPBS containing 1:1,000 diluted Goat anti-Rabbit IgG (H+L) Cross-Adsorbed Secondary Antibody, Alexa Fluor 488 or Alexa Fluor 647 (Catalog # A-11008 or A-21244, Thermo Fisher Scientific) and incubated for 30 min at room temperature. Cells were washed twice by using 200 µl of 0.1% Saponin in 3% BSA DPBS following by one time washing with 1 × DPBS. Cells were eventually resuspended in 100 µl of 1× DPBS and analyzed on the BD LSRII flow cytometry. Data were analyzed using the FlowJo software (Tree Star, Ashland, OR) and showed similar variance.

**Quantitative RT-PCR (qRT-PCR)**

Indicated ES cells were freshly collected, and total RNA was extracted by using Trizol reagent (15596026, Thermo Fisher Scientific) according to the manufacturer's instructions. RNA was then treated with RNase free DNase set (79254, Qiagen) and cDNA was synthesized using reagents supplied with the first strand cDNA synthesis kit (11483188001, Roche). Quantitative real-time PCR was performed using SYBR Green I (04707516001, Roche) on a LightCycler 480 instrument (Roche). Primer sequences are available upon request.

**Alkaline phosphatase (AP) staining**

Alkaline Phosphatase staining was performed using the Stemgent® Alkaline Phosphatase Staining Kit II (Stemgent, 00-0055) according to the manufacturer's instructions. In detail, $5 \times 10^5$ ES cells were seeded in a well of the 6-well plate a day before the experimental day. ES medium was aspirated, and cells were washed with 2 ml of 1 × PBST (1 × DPBS with 0.05% v/v of Tween20). Fix solution was added and incubated at room temperature for 2 minutes. Fix solution was then aspirated, and the fixed cells were washed with 2 ml of 1 × PBST. 1.5 ml of freshly prepared AP substrate solution was added in each well and incubated at room temperature for 12 minutes. Reaction was stopped by aspirating the AP Substrate Solution and the cells were washed twice with 2 ml of 1 × PBS. Cells were covered with 1 × PBS and directly used for imaging by using DMiL LED inverted tissue culture microscope (11521266, Leica) with Flexacam C1 camera. LAS X software was used for the data analysis.

**Seahorse metabolic assays**

Control or I-BETR CY2.4 albino mouse embryonic stem (ES) cells were seeded in a Seahorse XF96 Cell Culture Microplate witht the density of $5 \times 10^4$ cells/well. Cells were cultured in medium with or without glucose overnight before the assay. For measuring OCR and ECAR rates in the I-BETR ES cells, medium without I-BET was applied to the I-BETR cells for indicated length of time. Culture media were exchanged to Agilent Seahorse XF Base Medium (102353-100, Agilent) 1 hour prior to the assay. OCR rates were measured by Agilent Seahorse XF Cell Mito Stress Test Kit (103015-100, Agilent) under basal conditions and after the sequential addition of 1 µM Oligomycin, 0.5 µM FCCP (carbonyl cyanide p-(trifluromethoxy) phenylhydrazone), 0.5 µM Rotenone and Antimycin A for the final concentrations according to the manufacturer's instructions. ECAR rates were measured by Agilent Seahorse XF Glycolysis Stress Test Kit (103020-100, Agilent) under basal conditions and after the sequential addition of 0.5 µM Rotenone/Antimycin A and 50 mM 2-DG for the final concentrations according to the manufacturer's instructions. Both the OCR and ECAR assays were performed by using Agilent Seahorse XFe96 Analyzer. All values were calculated per well and normalized to cell number for each experiment. Each experiment was normalized to the average of all controls across all the experiments performed (n=3). The metrics of OCR assay were calculated according to the following equations: Basal Respiration = $OCR_{baseline} - Lowest\ OCR_{R+A}$ and Maximum Respiration = $Highest\ OCR_{FCCP} - Lowest\ OCR_{R+A}$. As proton efflux from live cells comprises both glycolytic and mitochondrial-derived acidification, for the ECAR assay, inhibition of mitochondrial function by Rotenone and Antimycin A enables calculation of mitochondrial-associated acidification. Subtraction of mitochondrial acidification to total proton efflux rate results in glycolytic proton efflux rate.

**RNAi-mediated knockdown of *Dusp4*, *Dusp6* and *Spry4***

ES cells are trypsinized, counted and diluted in ES media without antibiotic to a density of $2.5 \times 10^5$ cell/ml. $1 \times 10^5$ ES cells were seeded in a 6-well plate containing feeders and incubated in at 37°C with 5% $CO_2$ overnight. On the next day, mix 2.5 µl of 20 uM SMARTpool siRNA oligonucleotides (Dharmacon) targeted to mouse *Dusp4*, *Dusp6*, *Spry4* or a scrambled siRNA control (diluted in RNase-free buffer) with 197.5 µl of serum-free media in a microfuge tube (Tube 1) and incubate for 5 min at room temperature. In another tube, mix 4 µl of DharmaFECT® 1 with 196 µl of serum-free media (Tube 2), mix and incubate for 5 min at room temperature. Transfer the contents of Tube 1 into Tube 2, mix gently, and incubate for another 20 min at room temperature. Remove ES media without antibiotic from the wells and add 400 µl of transfection medium to each well. After a 30 min incubation at room temperature, top up the volume to 2ml for each well by using ES media without antibiotic (~1.6ml). Mock-transfected cells were given DharmaFECT 1 transfection reagent only. Incubate the cells in a $CO_2$ incubator for 48 hours and targeted gene expression was assessed by qRT-PCR. The order information for SMARTpool siRNA targeting of mouse *Dusp4*, *Dusp6* and *Spry4* are as follows: L-061306-00-0005, L-040050-00-0005 and L-059172-01-0005. The order information for mock non-targeting siRNA pool is D-001810-10.

**RNA-seq library preparation and analysis**

Generation and sequencing of RNA-seq samples

Three or four replicates were used for samples in RNA-seq assays. Freshly collected cells were used for total RNA extraction by using TRIzol reagent (Thermo Fisher Scientific) according to the manufacturer's instructions. Samples were either spiked-in or not with RNA Spike-In Mix (4456740, Thermo Fisher Scientific) following manufacturer's recommendations. Ribosomal and mitochondrial RNA was removed, and library preparation were performed by using 250 ng total RNA in all samples based on TruSeq Stranded Total RNA Ribo-Zero Gold library prep kit for Illumina (RS-122-2301, Illumina). Samples were sequenced at Genomics Resource Center, The Rockefeller University on Illumina NextSeq500.

Generation of Transcript Annotations

The Get Gene Annotation (GGA) pipeline (https://github.com/AdelmanLab/GetGeneAnnotation_GGA ; https://doi.org/10.5281/zenodo.5519927) was used to generate cell-type specific gene annotations using PRO-seq (control) and RNA-seq (+/- 1 μM Torin1) data generated in mESCs. Briefly 5' end of PRO-seq reads were used to call transcription start sites (TSSs) and assign a dominant TSS for each gene. RNA-seq data was then quantified by kallisto (version 0.45.1) and used to assign a transcript end site (TES) location for each gene. These consensus gene annotations were used for all subsequent analyses.

RNA-seq analysis

Raw RNA-seq reads were quality filtered (mean quality score ≥20) and first mapped to ERCC spike sequences using STAR (version 2.7.3a). Reads not mapping to spike-in sequences were aligned to the mm10 genome using the following parameters: --quantMode TranscriptomeSAM GeneCounts --outSAMtype BAM SortedByCoordinate --loutMultimapperOrder Random --outSAMattrIHstart 0 --outFilterType BySJout --outFilterMismatchNmax 2 --alignSJoverhangMin 8 --outSAMstrandField intronMotif --outFilterIntronMotifs RemoveNoncanonicalUnannotated --alignIntronMin 20 --alignIntronMax 1000000 --alignMatesGapMax 1000000 --outWigType bedGraph --outWigNorm None --outFilterScoreMinOverLread 0 --outFilterMatchNminOverLread 0. Duplicates were also removed using STAR. Gene counts were generated using the featureCounts function of the Rsubread package (version 2.0.1) using the GGA annotations described above. Per-gene normalized counts values were then quantified using DESeq2 (version 1.26.0) and differentially expressed genes identified. Given comparable spike-recovery values between samples, normalization was performed using DESeq2 generated size factors.

Ontology analysis:

For I-BETR RNA-seq comparisons, differentially expressed genes called by DESeq2 (Adjusted p-value < 0.001 and fold change > 2) were ranked by fold change. The top 225 upregulated genes were run through EnrichR (https://maayanlab.cloud/Enrichr/) and unique MSigDB Hallmark and

Reactome categories passing significance thresholds (p-val < 0.001). The same process was repeated for the 200 most downregulated genes ranked by fold change.

**TT-seq library preparation and analysis**

TT-seq library preparation:

CY2.4 albino B6 ES cells were plated in T175 tissue culture flasks one day prior to treatment. On the day of treatment, Torin1 was added to a final concentration of 1 μM to ES media. For control treatment, an equal volume of vehicle was added to media for the same length of treatment. 500 mM 4sU was added to the samples exactly at 10min to the end of the given length of treatments. Cells were rinsed with room-temperature PBS and harvested using trypsin and quenched with cold DMEM + 10% FBS. After an additional wash with PBS, cells were counted and resuspended in 1 mL Trizol per $2 \times 10^6$ cells. Samples were frozen immediately at -80°C until all the samples were ready for the downstream steps.

Prior to addition of chloroform to the lysates, samples were spiked-in with 5% cell counts based 4sU-labeled Drosophila S2 Trizol lysate (cells labeled with 4sU for 2 h and resuspended a concentration of 10 million cells/ml in Trizol). RNA was then isolated per the manufacturer's protocol. The aqueous phase was precipitated by addition of 2.5 volumes of 100% ethanol with 1.4mM DTT, incubation at -20°C for 2h. Pellets were collected by centrifugation at 20,000g for 30 min at 4°C and washed twice with 500 mL of 75% ethanol before resuspension in 180 mL of nuclease free water. Aliquots were removed for quantification by spectrophotometry and analysis of RNA integrity by Agilent TapeStation 4200 using RNA high sensitivity tapes. Samples with RIN > 9.5 were used for further processing.

Total RNA from previous step was treated to remove residual DNA by using RNeasy Micro Kit (Qiagen, 74004) combined with Amplification Grade DNase I (18068015, Invitrogen) and Superase-In RNase inhibitor (AM2694, Thermo Fisher Scientific). In detail, 40 µl of 1M DTT was added into 1mL of RLT buffer prior to the procedures. 200 µl of RNA sample was mix with 700 µl of RLT buffer and 500 µl of 100% ethanol. Samples were loaded onto a MinElute column and spun for 1min at 3,500g. 700 µl of RW1 was then added and spun the column for 2min at 14,000g. 5 µl of DNase I and 35 µl of RDD mixture was then added onto the column and incubate at room temperature for 30 min. After incubation, 660 µl of RW1 was added and column was spun for 1min at 14,000g. Wash the column with one time of 500 µl of RW1 buffer followed with one time of 500 µl of RPE buffer. Next, 500 µl of 80% ethanol was added and then spun the column for 2min at 14,000g. Repeat the last step once more and open the lid of the tubes for 5min at room temperature. 30 µl of RNase free water was added onto the column and incubate at room temperature for 2min and then spun at 200g for 2min. Add another 22 µl of RNase free water onto the column and repeat the same incubation following with spinning. Eventually the RNA samples were eluted by spun down for 2min at 14,000g and the concentration and quality of the samples were detected by Nanodrop and TapeStation respectively. Samples with RIN > 9.5 were used for further processing.

RNA samples were then chemically fragmented by addition of 20 mL cold 5 × fragmentation solution (375 mM Tris-HCl (pH 8.3), 562.5 mM KCl, 22.5 mM $MgCl_2$) and incubation at 94°C for 3.0 minutes. At the end of the fragmentation time, RNA was placed immediately on ice and 25 mL of cold 250 mM EDTA was added. RNA was precipitated by addition of 1/10 volume of 5 M NaCl, 2.5 volumes of 100% ethanol and incubation at -20°C overnight. RNA samples were pelleted, washed, quantified, and analyzed again as described above. The TapeStation results should show bulk of the signal at 200-1,000nt with the peak size around 450-600nt.

Fragmented RNA was biotinylated as described in Duffy et al. (2015) with the following modifications: the biotinylation reaction was performed in a total volume of 200 µl and allowed to incubate for 45 min in the dark. Excess biotin was removed using chloroform:isoamyl alcohol and Phase-Lock-Gel (Heavy/High density) tube (Qiagen, 129056) were used to separate organic and aqueous phases. Biotinylated RNA was resuspended in 100 µl of nuclease-free water and aliquots taken to use as the total RNA input fraction. In parallel, Dynabeads MyOne Streptavidin C1 (Invitrogen, 65001) were prepared for binding to render them RNase-free: for each sample, 25 µl of beads were used and treated in batch to render them RNase free. The beads were incubated 10 min in the decon solution (100 mM NaOH, 50 mM NaCl) and placed on a magnetic stand. Then the beads were washed as follows: resuspending the beads fully for each wash, twice with 500 µl of 100 mM NaCl, twice with 500 µl of 1 × TTseq wash solution (100 mM Tris-HCl (pH 7.4), 10mM EDTA, 1 M NaCl, 0.05% Tween 20 in nuclease free water to which 1 µl SuperaseIN RNase Inhibitor (AM2694, Thermo Fisher Scientific) per 1mL solution is added prior to use), once in 500 µl of 0.3 × TTseq wash solution, and finally resuspended in 52 µl /sample of 0.3 × TTseq wash solution and 1 µl/sample Superase-In RNase inhibitor (Invitrogen, AM2696). Biotinylated RNA was heated at 65°C for 5min, placed on ice for 2 min, and mixed with 50 µl of prepared beads. Samples were rotated at room temperature in the dark for 30 min. After binding, the tubes were placed on a magnetic rack and the beads were washed 4 times with 500 µl of 1 × TT-seq wash solution to remove unbound RNA, fully resuspending for each wash. The wash solution was removed, and the beads resuspended in 50 µl of 100mM of DTT (freshly diluted from 1 M DTT stock) and incubated in the dark for 15 min at room temperature. The eluted RNA was recovered, and the elution step repeated with an additional 50 µl of 100mM of DTT.

The combined eluates were purified using Norgen RNA clean-up and concentration MicroElute kit (Norgen #61000) following the manufacturer's instructions for small RNA enriched samples. Final elution was performed in 14 µl of nuclease-free water and the eluate was reapplied to the column for a total of two elution steps. A Qubit RNA high sensitivity reagent kit was used to quantify the input RNA and enriched RNA. Yields of 1-2% were typical. 340-400 ng of enriched RNA was used for library construction with the TruSeq RNA Library Preparation Kit v2, Set A and TruSeq RNA Library Preparation Kit v2, Set B (Illumina, RS-122-2001 and RS-122-2002) for individually indexed the samples. After 5 cycles of PCR, samples were removed from the thermal cycler and a test PCR was performed to determine the optimal number of final cycles. Libraries were pooled and pair-end sequenced on a NovaSeq SP: 2 × 50 flowcell.

<u>TT-seq analysis</u>:

TT-seq reads were filtered (mean quality score ≥20) and mapped to the *Drosophila* (dm6) genome using STAR (version 2.7.3a). Reads not mapping to spike sequences were aligned to the mm10 using the following parameters: --quantMode TranscriptomeSAM GeneCounts --outSAMtype BAM SortedByCoordinate --loutMultimapperOrder Random --outSAMattrIHstart 0 --

outFilterType BySJout --outFilterMismatchNmax 2 --alignSJoverhangMin 8 --outSAMstrandField intronMotif --outFilterIntronMotifs RemoveNoncanonicalUnannotated --alignIntronMin 20 --alignIntronMax 1000000 --alignMatesGapMax 1000000 --outWigType bedGraph --outWigNorm None --outFilterScoreMinOverLread 0 --outFilterMatchNminOverLread 0. Duplicates were also removed using STAR. Gene counts were generated using the featureCounts function of the Rsubread package (version 2.0.1) using the same GGA annotation used for RNA-seq. Differentially expressed genes and per-gene normalized counts values were then quantified using DESeq2 (version 1.26.0). Given comparable spike-recovery values between samples, normalization was performed using DESeq2 generated size factors. These normalization factors were also used to generate combined bedGraphs using coverage from all replicates across each condition.

Cluster analysis:

For clustering of TT-seq data, protein-coding genes that were differentially expressed (Adjusted p-value < 0.005 and fold change > 1.5) at any time point of Torin-1 treatment were considered (1,065 genes). The clara function of the cluster package (R, version 2.1.4) was used to calculate dissimilarity between genes, resulting in 3 distinct clusters. Genes were represented in their respective clusters using the $\log_2$FoldChange values assigned via DESeq2.

Ontology analysis:

For TT-seq data, genes from each cluster (1-3) were independently run through EnrichR to extract MSigDB Hallmark ontologies. Significant categories (p-adj < 0.05) were represented for each corresponding cluster set.

**ChIP-seq library preparation and analysis**

ChIP-seq library preparation:

CY2.4 ES cells were plated at a density of $2.5 \times 10^5$ cells/ml in ES medium (as described before) and cultured for 24 hours. (Cells were always grown on MEFs except one passage on gelatinized tissue culture plates before the experiment). ES cells were trypsinized and resuspended in a concentration of $1 \times 10^6$ cells/ml in DPBS (room temperature). $2 \times 10^7$ cells were used for each ChIP sample. Pierce 16% formaldehyde (28906, Thermo Fisher Scientific) was added to a final concentration of 1% and cross-linking was performed for 10 minutes at room temperature. Cross-linking was then terminated by adding 2.5M glycine to a final concentration of 0.125M at room temperature. Cells were washed with cold DPBS twice and pelleted by spun down at 1,260g for 6 minutes at 4°C. Cells were resuspended in LB1 buffer (50mM Hepes-KOH, pH7.9, 140mM NaCl, 1mM EDTA, 10% glycerol, 0.5% NP40, 1% Triton X-100, 1 × protease inhibitor, 1 × phosphatase inhibitor cocktail set I and 100μM TSA) and incubate for 20 minutes rotating at 4°C. Cells were pelleted for 6 minutes at 1,350 g at 4°C. Cells were then resuspended in LB2 buffer (10 mM Tris pH 8.0, 200 mM NaCl, 1 mM EDTA, 0.5 mM EGTA, 1 × protease inhibitor, 1 × phosphatase inhibitor cocktail set I and 100μM TSA) and incubated for 5 minutes rotating at 4°C. Cells were pelleted for 6 minutes at 1,350 g at 4°C and following by resuspended in LB3 (10 mM Tris pH 8.0, 100 mM NaCl, 1 mM EDTA, 0.5 mM EGTA, 0.1% sodium-deoxycholate, 0.5% sodium lauroyl sarcosinate, 1% Triton X-100, 1 × protease inhibitor, 1 × phosphatase inhibitor cocktail set I and 100µM TSA). Cells were sonicated using Focused-Ultra sonicator (E220, Covaris) for 12 minutes to generate chromatin fragments range from 200 to 600 bp of DNA in length according to the manufacturer's instructions. Fragmented chromatin samples were then spun down at 20,000g for 30min at 4°C. Dynabeads (Invitrogen Dynabeads anti-rabbit M-280) were pre-blocked with 0.5% BSA in 1 × DPBS and were incubated with 10 µg of CIC (PA1-46018, Thermo Fisher Scientific) antibodies for 8 hours. Dynabeads were washed twice by using 0.5% BSA in 1 × DPBS and fragmented chromatin was added to antibody-bead complex and incubated overnight by rotating at 4°C.

Beads were sequentially washed three times with wash buffer 1 (50mM Hepes pH7.5, 500mM NaCl, 1mM EDTA, 1mM EGTA, 1% Triton, 0.1% NaDoc, 0.1% SDS) and three times with wash buffer 2 (20mM Tris pH 8, 1mM EDTA, 250mM LiCl, 0.5% NP40, 0.5% NaDoc) at 4°C, followed by washing one time with 1 × TE buffer at room temperature. Pulled-down chromatin fragments was eluted by adding elution buffer (50 mM, Tris pH 8.0, 10 mM EDTA, 1% sodium dodecyl sulfate, 20ug/ml RNaseA) to the beads and incubated with shaking at 65°C for 30 minutes. Beads were then collected by spun down briefly and the supernatants were transferred into a new tube. Reversal of crosslinking was performed at 65°C overnight in a thermocycler. 2 µl of DNase-free RNase (11119915001, Roche) was added and incubated for 1 hour at 37°C following with 2 µl of Proteinase K treatment for 2 hours at 55°C. DNA was purified by using Qiagen PCR purification kit and resuspended in 10mM Tris-HCL.

The concentrations of the pulled-down DNA fragments were measured by Qubit dsDNA HS and BR Assay Kits (Q32851, Invitrogen) and ChIP libraries were prepared by using NEBNext Ultra II DNA Library Prep Kit for Illumina (E7103S, NEB) with size selection step. The concentration and size distribution of the libraries were then measured by Qubit dsDNA HS and BR Assay Kits and tapestation respectively. 100nM of each library was pooled together with a maximal 12 samples in a line for Illumina NextSeq500 sequencing in the Genomics Resource Center of the Rockefeller University.

ChIP-seq analysis:

For CIC ChIP-seq samples, raw reads were filtered (mean quality score ≥20) and adapter sequences were removed using cutadapt (version 1.14). Reads were then mapped to the mm10 reference genome using bowtie (version 1.2.2). Fragment midpoints were approximated by shifting coordinates downstream by 75bp and were converted to bedGraphs with a bin size of 50bp using the script bowtie2stdBedGraph.pl (https://github.com/AdelmanLab/NIH_scripts; https://doi.org/10.5281/zenodo.5519915). BedGraphs were then normalized by total sequencing depth and converted to bigWigs for visualization on the UCSC genome browser.

**PRO-seq library preparation and analysis**

PRO-seq library preparation

Control mESCs were permeabilized and subjected to PRO-seq library preparation as previously described(*2*). Final libraries were pooled and sequenced paired end on an Illumina NovaSeq platform.

PRO-seq analyses

For mapping PRO-seq libraries, the following pipeline was used. Dual unique molecular identifiers (UMIs) 6nt in length were extracted using UMI tools (https://doi.org/10.1101/gr.209601.116). Read pairs were trimmed using cutadapt (version 1.14). An additional nucleotide was removed from read 1 (3' end read) using seqtk (https://github.com/lh3/seqtk) to ensure non-overlapping read pairs in mapping. Reads were mapped to a combined genome comprised of both spike (dm6) and reference (mm10) genomes using bowtie2, which were then separated based upon respective genome. Using previously identified UMIs, mapped reads were deduplicated, then sorted using samtools (version 1.3.1). Reference reads were then separated into 5' and 3' end reads and subsequently converted to strand-specific bedGraph format using the script bowtie2stdBedGraph.pl (https://github.com/AdelmanLab/NIH_scripts; https://doi.org/10.5281/zenodo.5519915). Successfully mapped 5' bedGraphs were used for determining gene annotation information.

**Accession numbers**

The data discussed in this publication have been deposited in NCBI's Gene Expression Omnibus (GEO) and are accessible through GEO Super Series accession number GSE249392.

**Statistical analysis**

All the data were processed using Prism 9. For analysis of the correlation of gene expression between 160 ribosomal and all the genes, the nonparametric Mann–Whitney U test with null hypothesis for randomly selected values X and Y from two populations was used. Statistical significance is indicated by *, $p < 0.05$; **, $p < 0.01$; ***, $p < 0.001$; and ****, $p < 0.0001$. For all the other assays, Unpaired t-test was employed and performed for the analysis. Statistical significance is indicated by *, $p < 0.05$; **, $p < 0.01$; ***, $p < 0.001$; and ****, $p < 0.0001$.

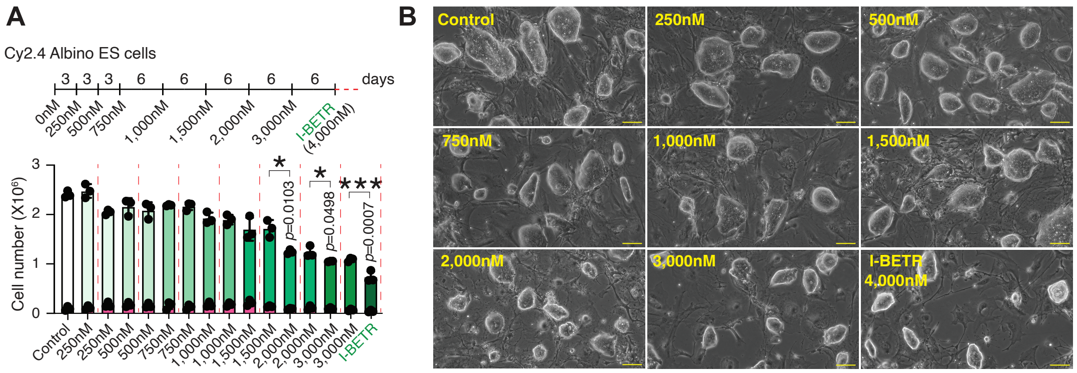


**Fig. S1. Generation of I-BET151 resistant (I-BETR) ES cells. (A)**, Stepwise generation of I-BETR murine ES cells. The bar graph shows the ES cell number at the indicated concentration of I-BET at each step of ES cell treatment. Number of dead cells is highlighted in pink. *, $p$<0.05, ***, $p$<0.001, Unpaired t-test had been used for the definition of significant slowing down of the cell growth at the indicated I-BET concentration as compared to the previous concentration. **(B)**, Progressive increase in I-BET concentration does not affect the gross morphology and appearance of ES cell colonies. The indicated concentration of I-BET151 is shown. Scale bar, 100µm.
